# Enhanced antitumour activity of combined PD-1 blockade and HER2-targeted therapy is associated with reduced epithelial-mesenchymal transition and extracellular matrix-remodelling signatures in gastric cancer

**DOI:** 10.64898/2026.11.08.765160

**Authors:** Peter Kok-Ting Wan, Shuang Peng, Xiangrui Zhang, Jian Wang

## Abstract

**Background:** The combination of PD-1 blockade, HER2-targeted therapy and chemotherapy improves outcomes in HER2-positive advanced gastric and gastro-oesophageal junction adenocarcinoma. However, chemotherapy can independently alter the tumour microenvironment, making the biological effects of PD-1 and HER2 targeting difficult to distinguish. We therefore investigated how PD-1 and HER2 blockade, alone and in combination, affect tumour growth and associated transcriptional programmes.

**Methods:** Immunodeficient NOG mice received human PBMCs by tail-vein injection. Three days later, NCI-N87 gastric cancer cells were implanted subcutaneously. Mice were randomised to vehicle, pembrolizumab, trastuzumab, or combination treatment. Tumour growth was assessed, and bulk RNA sequencing was used to identify treatment-associated changes in gene expression, pathway enrichment, and tumour, stromal, and immune cell-associated signatures.

**Results:** Combined treatment produced greater tumour growth control than either monotherapy. Trastuzumab-treated tumours showed enrichment of immune activation, epithelial-mesenchymal transition (EMT), and TGFβ-associated signatures, whereas pembrolizumab alone produced a limited transcriptional effect. Combination treatment was associated with reduced proliferation, EMT, and stromal-remodelling signatures, alongside increased cytotoxic immune-associated signatures. Expression of extracellular matrix-associated genes, including VCAN, SPARC, POSTN, collagen genes, and FAP, was lower in combination-treated tumours. Complement-related transcriptional pathways were also enriched.

**Conclusions:** In this chemotherapy-free model, combined PD-1 blockade and HER2-targeted therapy was associated with greater tumour growth control, enrichment of cytotoxic immune-associated transcriptional signatures, and reduced EMT and extracellular matrix remodelling signatures. These findings provide hypotheses for further mechanistic validation.

**Mini-abstract:** In chemotherapy-free humanised mice, pembrolizumab plus trastuzumab was associated with stronger tumour control, increased cytotoxic immune signatures, and reduced proliferation, EMT, and extracellular matrix remodelling programmes.

## Introduction

Advanced gastric and gastro-oesophageal junction adenocarcinoma remains therapeutically challenging, particularly in patients with unresectable or metastatic disease. HER2 overexpression or amplification identifies a clinically actionable subgroup, with HER2 positivity reported in approximately 20% of gastric and gastro-oesophageal junction cancers^1^. The clinical value of HER2 targeting was established by the ToGA trial, in which adding trastuzumab to platinum and fluoropyrimidine-based chemotherapy improved median overall survival from 11.1 to 13.8 months^2^. However, trastuzumab is not typically used as a standalone therapy in gastric cancer, and responses to trastuzumab-containing regimens remain variable and frequently non-durable^3–5^. These limitations highlight the need to better understand how HER2-targeted therapy interacts with the tumour microenvironment and how it may be optimally combined with immunotherapy.

Trastuzumab exerts anti-tumour activity not only through HER2 receptor blockade, but also through Fc-dependent immune effector mechanisms, including antibody-dependent cellular cytotoxicity mediated by Fc receptor-bearing immune cells such as NK cells^6,7^. This immune component provides a strong rationale for combination with immune checkpoint blockade, as trastuzumab-mediated tumour cell killing may promote antigen release, dendritic-cell cross-presentation, and downstream T cell priming^8,9^. Pembrolizumab, an anti-PD-1 antibody, may therefore enhance and sustain anti-tumour T cell responses that arise indirectly from trastuzumab-mediated immune engagement. In gastric cancer models, trastuzumab has also been reported to induce PD-L1 upregulation through NK-cell-derived IFN-γ, suggesting that HER2-targeted therapy can generate a checkpoint response that may be counteracted by PD-1 blockade^10,11^.

The clinical benefit of this combination has been demonstrated in the phase III KEYNOTE-811 trial, in which the addition of pembrolizumab to trastuzumab and fluoropyrimidine/platinum chemotherapy improved objective response rate, progression-free survival, and overall survival in previously untreated HER2-positive advanced gastric or gastro-oesophageal junction adenocarcinoma. In the PD-L1 CPS ≥1 population, the addition of pembrolizumab improved median progression-free survival from 7.3 to 10.9 months, and median overall survival from 15.7 to 20.0 months^12^. These findings led to regulatory approval of pembrolizumab in combination with trastuzumab and chemotherapy as a first-line treatment for HER2-positive, PD-L1-positive advanced gastric or gastro-oesophageal junction adenocarcinoma^13^.

Despite these clinical advances, the biological basis of therapeutic cooperation between PD-1 blockade and HER2-targeted therapy remains incompletely defined. In the clinical setting, pembrolizumab is administered with trastuzumab and cytotoxic chemotherapy, and chemotherapy can independently modulate anti-tumour immunity through antigen release, inflammatory signalling, immune cell recruitment, cell death and stromal remodelling^14–16^. Therefore, although KEYNOTE-811 establishes the clinical benefit of adding pembrolizumab to trastuzumab-based chemotherapy, it does not fully resolve the direct interaction between PD-1 blockade and HER2-targeted therapy. A chemotherapy-free experimental setting provides a useful mechanistic window to examine how pembrolizumab and trastuzumab interact within the gastric cancer tumour microenvironment. In particular, the transcriptional programmes that distinguish monotherapy effects from combination-driven tumour growth control in gastric cancer remain insufficiently resolved.

In this study, we investigated pembrolizumab, trastuzumab and their combination in gastric cancer xenografts established in PBMC-engrafted humanised mice. This model enabled controlled *in vivo* comparison of PD-1 blockade, HER2-targeted therapy and dual treatment in the absence of chemotherapy. We observed enhanced control on tumour growth following combination treatment and performed bulk RNA sequencing to examine treatment-associated changes in the tumour microenvironment. By comparing the transcriptional profiles associated with pembrolizumab, trastuzumab monotherapy and their combination, this study identifies signatures associated with the enhanced anti-tumour activity observed with dual PD-1 and HER2 targeting.

## Results

### Combination of pembrolizumab and trastuzumab enhances tumour growth control

To evaluate the therapeutic potential of PD-1 blockade and HER2-targeted therapy *in vivo*, we implanted the human gastric cancer cell line NCI-N87 into PBMC-engrafted humanised NOG mice. Tumour-bearing mice were treated with PBS, pembrolizumab, trastuzumab, or the combination of pembrolizumab and trastuzumab (Figure 1A). Tumours in the PBS control group showed progressive growth throughout the study. Both monotherapies reduced tumour growth compared with PBS, with trastuzumab showing a marginally greater reduction in tumour volume at later time points (Figure 1B).

**Figure 1.**
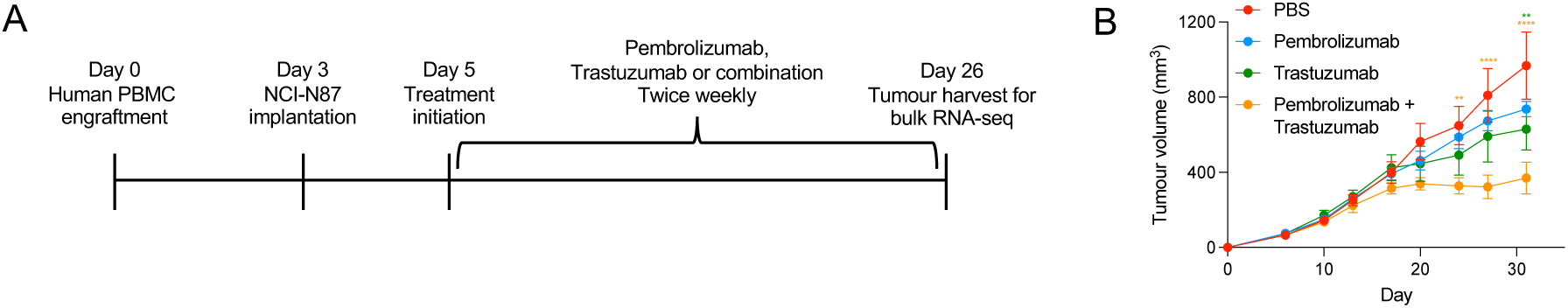
Combined pembrolizumab and trastuzumab treatment enhances anti-tumour activity in PBMC-engrafted gastric cancer xenograft model. Gastric cancer xenografts established in PBMC-engrafted humanised mice were treated with PBS, pembrolizumab, trastuzumab, or pembrolizumab plus trastuzumab. (A) Schematic overview of the experimental design, including PBMC engraftment, NCI-N87 tumour implantation, treatment initiation and tumour harvest for bulk RNA-seq. (B) Longitudinal tumour-growth curves showing tumour volume over time in each treatment group. Data are shown as mean ± SEM.

The combination of pembrolizumab and trastuzumab produced more pronounced tumour growth control than either monotherapy. Tumour growth in the combination group stabilised from approximately day 17 onward, and tumours remained substantially smaller than those in the PBS and monotherapy groups through the end of the study. These findings indicate that combined PD-1 blockade and HER2-targeted therapy produced a stronger anti-tumour effect than either treatment alone.

### Transcriptional profiling of monotherapy-treated tumours reveals distinct effects of pembrolizumab and trastuzumab

We next performed bulk RNA-seq analysis to define how each monotherapy altered the transcriptional state of gastric cancer xenografts. Following filtering and TMM normalisation, sample-level quality control was performed using principal-component analysis, sample-sample correlation, detected-gene counts and logCPM distribution plots (Supplementary Figure 1A-D). These analyses showed broadly consistent sequencing depth and expression distributions across samples, supporting the suitability of the dataset for downstream differential expression analysis.

Compared with PBS-treated controls, pembrolizumab monotherapy induced a relatively limited transcriptional shift, with 163 differentially expressed genes identified, comprising 57 upregulated and 106 downregulated genes (Figure 2A). Notable differentially expressed genes included transcripts associated with lipid metabolism (FADS2, SCD), tumour pH regulation (CA9) and epithelial differentiation (KRT13). In contrast, trastuzumab induced broader differential gene expression, with 474 differentially expressed genes, including 219 upregulated and 255 downregulated genes (Figure 2B). Upregulated genes included immune cell recruitment and activation-associated genes, such as CCL2, CCL22, CCR4 and IL2RA, suggesting potential recruitment of monocytes/macrophages and CCR4^+^ T cells, including regulatory T cells, within the tumour microenvironment. Heatmap analysis of the top 15 differentially expressed genes from each comparison further highlighted the distinct treatment-associated transcriptional patterns induced by pembrolizumab and trastuzumab (Figure 2C, Supplementary Figure 2).

**Figure 2.**
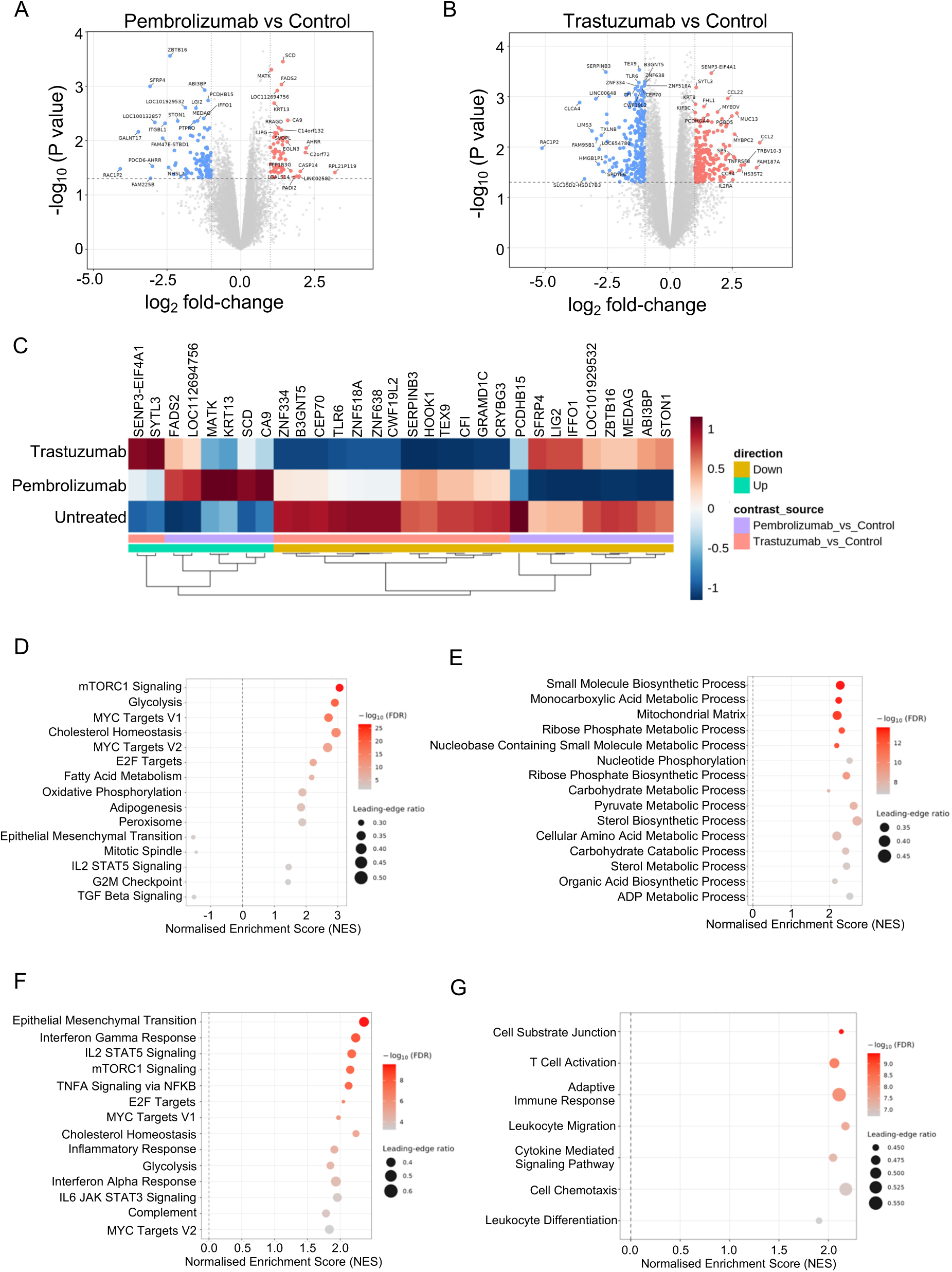
Pembrolizumab and trastuzumab monotherapies induce distinct transcriptional programmes in gastric cancer xenografts. Bulk RNA-seq was performed on gastric cancer xenografts from PBMC-engrafted humanised mice treated with PBS, pembrolizumab or trastuzumab. (A, B) Volcano plots showing differentially expressed genes in pembrolizumab-treated (A) and trastuzumab-treated (B) tumours compared with PBS controls, respectively. Differentially expressed genes were defined using *P* < 0.05 and absolute log2 fold-change ≥1. (C) Heatmap showing top differentially expressed genes from the pembrolizumab-versus-control and trastuzumab-versus-control comparisons. (D, E) Hallmark (D) and C5 Gene Ontology (E) GSEA of pembrolizumab-treated tumours versus PBS controls. (F, G) Hallmark (F) and C5 Gene Ontology (E) GSEA of trastuzumab-treated tumours versus PBS controls. For GSEA plots, positive normalised enrichment scores indicate enrichment in the treatment group relative to PBS controls, whereas negative scores indicate enrichment in PBS controls. Dot colour represents -log10(FDR), and dot size represents the leading-edge ratio.

Pathway analysis by Hallmark GSEA showed enrichment of mTORC1 signalling, glycolysis, MYC targets, cholesterol homeostasis, E2F targets, fatty acid metabolism and oxidative phosphorylation following pembrolizumab monotherapy (Figure 2D, Supplementary Figure 3A). Rather than indicating tumour-intrinsic proliferation alone, these metabolic and growth-associated programmes may reflect increased biosynthetic and energetic demand within the tumour microenvironment following PD-1 blockade. As pembrolizumab acts by relieving PD-1-mediated inhibitory signalling, its transcriptional effect may be more associated with sustaining the proliferation, survival and effector function of pre-existing or antigen-engaged T cells, rather than inducing a broad *de novo* T cell activation programme. This interpretation is further supported by enrichment of IL2-STAT5 signalling and C5 Gene Ontology terms related to small molecule biosynthetic processes, mitochondrial matrix, nucleotide metabolism and ribose phosphate metabolism (Figure 2E, Supplementary Figure 3B), which are consistent with the metabolic requirements of sustained lymphocyte expansion and cytotoxic activity. Surprisingly, trastuzumab treatment showed strong positive enrichment of epithelial-mesenchymal transition (EMT), despite the observed reduction in tumour growth *in vivo* (Figure 2F, Supplementary Figure 3C). Together with enrichment of IL6-JAK-STAT3 signalling and the C5 Gene Ontology term cell substrate junction, this suggests that trastuzumab may induce tumour adaptation programmes associated with EMT and junctional remodelling (Figure 2G, Supplementary Figure 3D).

Trastuzumab also showed positive enrichment of immune-associated Hallmark pathways, including IFNγ response, TNFα signalling via NF-κB, inflammatory response, complement and IL2-STAT5 signalling (Figure 2F). Consistently, C5 Gene Ontology analysis showed enrichment of T cell activation, adaptive immune response, leukocyte migration, cytokine-mediated signalling, cell chemotaxis and leukocyte differentiation (Figure 2G). However, because leukocyte migration and chemotaxis may reflect recruitment of either cytotoxic lymphocytes or immunosuppressive lymphoid/myeloid populations, further analysis is required to define the specific immune cell populations involved. Overall, trastuzumab monotherapy appears to induce both immune activation and EMT/matrix-remodelling-associated programmes, indicating a complex response in the tumour microenvironment.

### Combination treatment is associated with reduced mesenchymal- and tumour growth-associated signatures

To define how combination of pembrolizumab and trastuzumab altered broader biological states within the tumour microenvironment compared with monotherapy, we performed ssGSEA across curated tumour-, stromal- and metabolism-associated signatures. Combination treatment reduced epithelial-mesenchymal transition (EMT), TGFβ signalling and mesenchymal identity signatures (Figure 3A). This was notable because untreated control tumours showed enrichment of mesenchymal identity than tumour epithelial identity, while trastuzumab monotherapy further increased the mesenchymal programme despite its anti-tumour activity. By contrast, the combination showed lower mesenchymal and higher epithelial-associated signature scores. These findings suggest that trastuzumab monotherapy may induce compensatory mesenchymal or tissue-remodelling adaptation, whereas the addition of PD-1 blockade was associated with attenuation of this transcriptional signature. Given the association of EMT, TGFβ signalling and mesenchymal remodelling with tumour plasticity and therapeutic resistance, their reduction in the combination group provides a plausible transcriptional basis for the enhanced anti-tumour activity observed with combination treatment.

**Figure 3.**
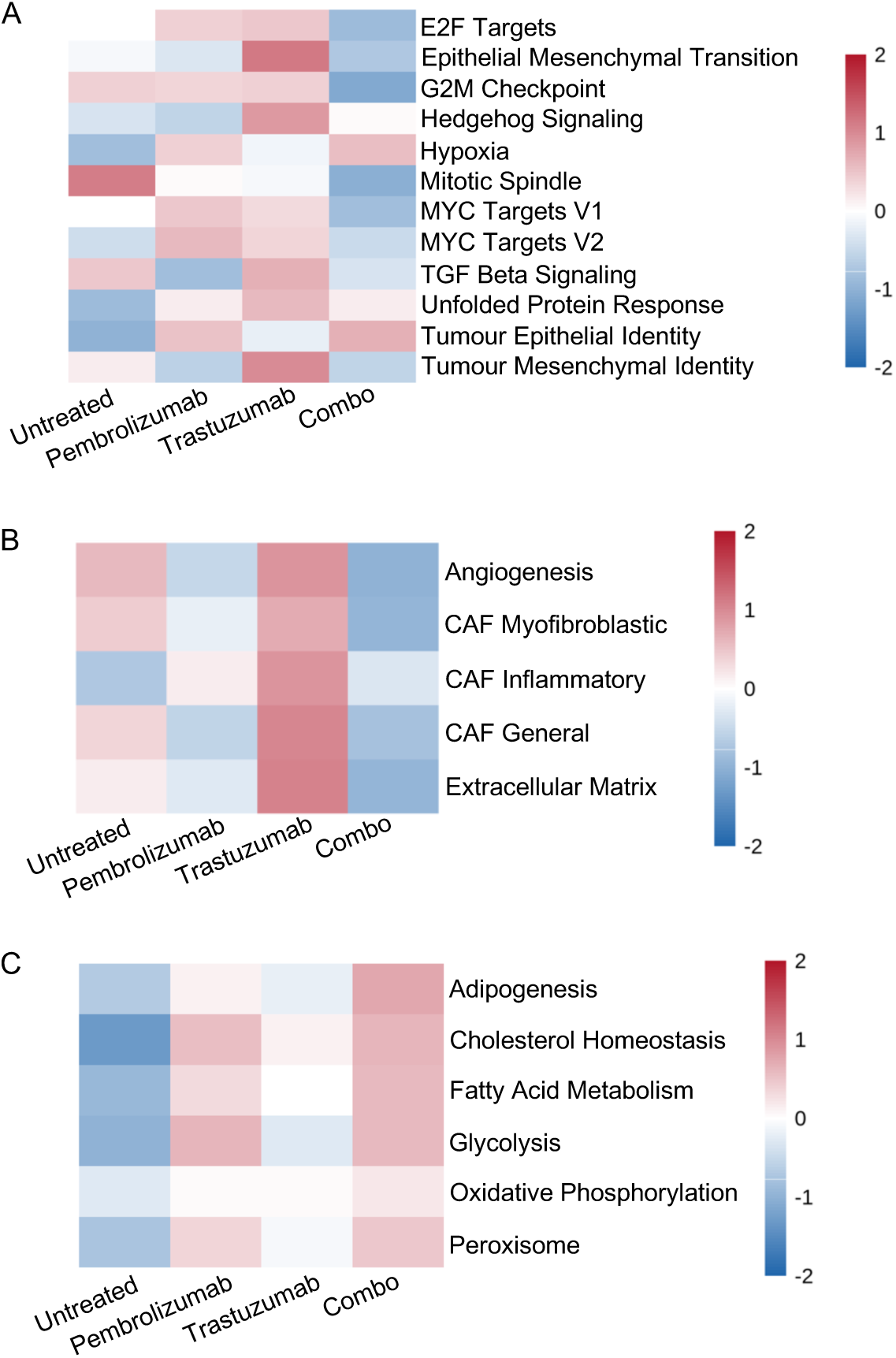
Combination treatment suppresses proliferative, mesenchymal and stromal programmes. ssGSEA was performed on bulk RNA-seq data from gastric cancer xenografts treated with PBS, pembrolizumab, trastuzumab, or pembrolizumab plus trastuzumab. Heatmap of tumour-associated pathways (A), stromal-associated signatures (B) and metabolic pathways (C). Heatmap colours represent mean scaled ssGSEA scores for each treatment group, expressed as z-scores.

Combination-treated tumours also showed reduced activity of multiple growth-associated programmes, including E2F targets, G2M checkpoint, mitotic spindle and MYC targets, compared with untreated or monotherapy-treated tumours (Figure 3A). Because these signatures can reflect proliferative activity across both tumour and immune compartments in bulk RNA-seq, they should be interpreted cautiously. However, in the context of the stronger tumour growth inhibition observed *in vivo*, their reduction likely reflect the suppression of overall tumour proliferation.

### Combination treatment is associated with lower stromal-remodelling signatures

Stromal signature analysis showed that untreated control tumours were enriched for angiogenesis, myofibroblastic CAF, general CAF and extracellular matrix (ECM) programmes, indicating a stroma-rich baseline tumour microenvironment (Figure 3B). Pembrolizumab monotherapy appeared to reduce these stromal-remodelling signatures while selectively increasing the inflammatory CAF signature. In contrast, trastuzumab monotherapy enriched all the tested stromal-remodelling programmes, including angiogenesis, CAF signatures, and ECM pathways. Combination-treated tumours showed reduced activity of these stromal-associated signatures compared with monotherapy-treated tumours and untreated controls, suggesting that dual PD-1 and HER2 blockade attenuates stromal-remodelling programmes that are present at baseline or further enhanced by trastuzumab monotherapy.

Metabolic pathway analysis showed that the combination-treated tumours retained enrichment of lipid- and glycolysis-related programmes, including adipogenesis, cholesterol homeostasis, fatty acid metabolism, glycolysis and peroxisome signatures, while oxidative phosphorylation remained comparatively unchanged (Figure 3C). Notably, this metabolic profile more closely mirrored pembrolizumab than trastuzumab monotherapy, suggesting that the metabolic component of the combination response may be primarily driven by PD-1 blockade, which may reflect sustained activation and expansion of immune cells, particularly T cells, as shown in Figure 2. However, because these signatures were derived from bulk RNA-seq, they likely reflect integrated metabolic activity across tumour, stromal and immune compartments.

### Exploratory deconvolution identifies treatment-associated immune cell signatures

To examine how each treatment altered the immune landscape, we applied xCell deconvolution to the bulk RNA-seq data (Figure 4). Pembrolizumab-treated tumours showed enrichment of naïve CD8^+^ T cell and common lymphoid progenitor-associated signatures, but showed limited enrichment of cytotoxic effector signatures. This suggests that PD-1 blockade alone was associated with relatively modest changes in cytotoxic immune-associated transcriptional programmes.

**Figure 4.**
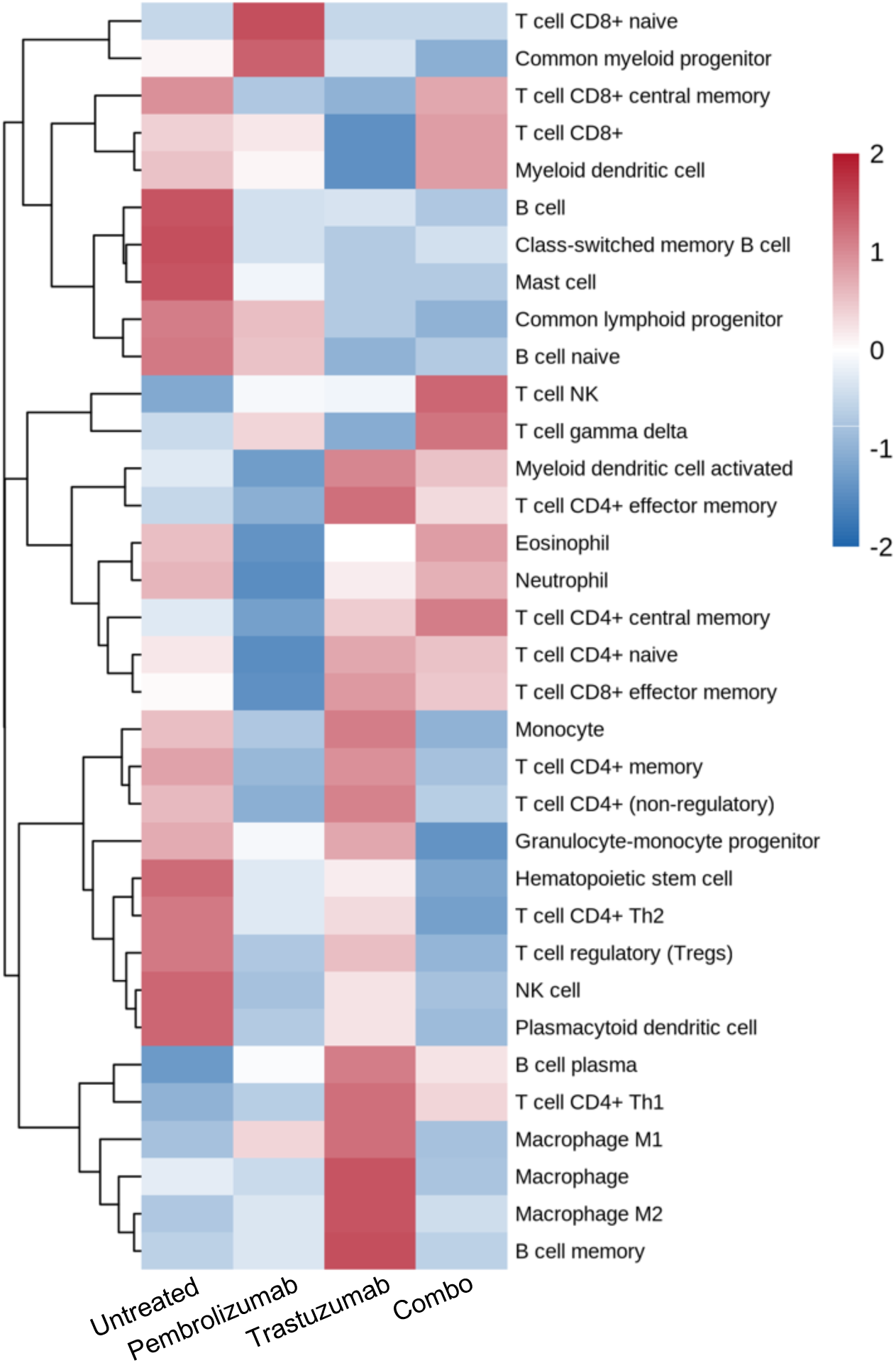
Combination treatment reshapes the immune landscape. Immune cell enrichment analysis was performed on bulk RNA-seq data from gastric cancer xenografts treated with PBS, pembrolizumab, trastuzumab, or pembrolizumab plus trastuzumab. Heatmap colours represent mean z-scored immune cell enrichment scores across treatment groups.

In contrast, trastuzumab-treated tumours showed strong enrichment of regulatory T cell, monocyte, macrophage and granulocyte-monocyte progenitor-associated signatures (Figure 4). The simultaneous enrichment of both M1-like and M2-like macrophage signatures suggests a mixed myeloid response involving inflammation and tissue-remodelling. This is consistent with the earlier observation that trastuzumab increased stromal, TGFβ and EMT-associated programmes.

Combination-treated tumours showed enrichment of multiple cytotoxic or effector lymphocyte populations, including CD8^+^ T cells, memory T cells, NK T cells and γδ T cells, together with dendritic cell-associated signatures while reducing the relative dominance of the macrophages and regulatory T cell observed in trastuzumab monotherapy. (Figure 4). Because xCell infers cell type-associated enrichment from transcriptional profiles rather than directly measuring immune cell abundance, these results were interpreted as exploratory immune cell-associated signatures.

### Combination treatment suppresses monotherapy-associated resistance programmes

To identify transcriptional programmes that may underlie the synergistic response observed *in vivo*, we directly compared pembrolizumab plus trastuzumab against each monotherapy. This analysis allowed us to distinguish pathways preferentially altered by the combined treatment from those induced by either agent alone.

In the comparison between combination treatment and pembrolizumab monotherapy, the volcano plot highlighted several immune-associated genes preferentially increased in the combination group, including the checkpoint receptor LAG3, T cell receptor genes TRBV9, TRBV4-2 and TRAV13-1, the innate-like T cell-associated transcription factor ZBTB16, and the lymphocyte-recruiting and tertiary lymphoid structure-associated chemokine CXCL13 (Figure 5A, Supplementary Figure 4). In contrast, genes linked to mitotic regulation, including KIF18A and KIF4A, were reduced in the combination group.

**Figure 5.**
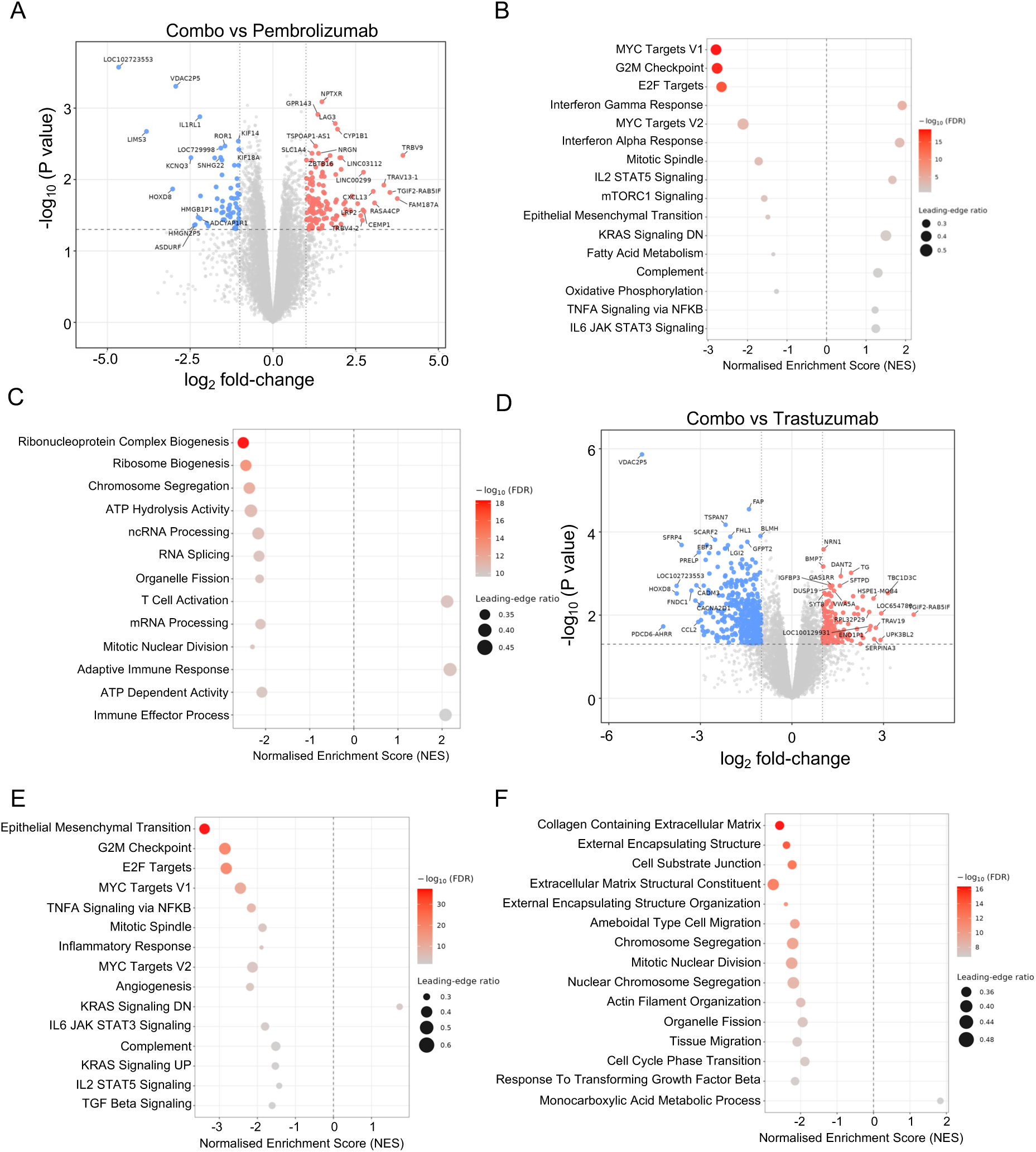
Combination treatment suppresses proliferative, mesenchymal and extracellular matrix-remodelling programmes relative to monotherapy. Bulk RNA-seq analysis was performed to compare pembrolizumab plus trastuzumab with each monotherapy. (A) Volcano plot showing differentially expressed genes in combination-treated tumours compared with pembrolizumab-treated tumours. Hallmark (B) and C5 Gene Ontology (C) GSEA comparing combination treatment with pembrolizumab. (D)Volcano plot showing differentially expressed genes in combination-treated tumours compared with trastuzumab-treated tumours. Hallmark (E) and C5 Gene Ontology (F) GSEA comparing combination treatment with trastuzumab. Positive NES indicates enrichment in the combination group, whereas negative NES indicates enrichment in the corresponding monotherapy group. Dot colour represents -log10(FDR), and dot size represents the leading-edge ratio.

Pathway analysis showed that combination treatment was associated with negative enrichment of proliferative and cell-cycle programmes, including G2M checkpoint, E2F targets and MYC targets (Figure 5B, Supplementary Figure 5A). C5 Gene Ontology analysis similarly showed reduced chromosome segregation, mitotic nuclear division, DNA replication-associated and ribonucleoprotein-related processes in the combination group (Figure 5C, Supplementary Figure 5B). In contrast, immune-related pathways, including IFNα response, IFNγ response, IL2-STAT5 signalling, T cell activation and adaptive immune response were relatively enriched (Figure 5B, C). These findings suggest that adding trastuzumab to pembrolizumab suppresses tumour growth programmes while enhancing immune-associated signalling.

Compared with trastuzumab monotherapy, combination treatment showed reduced expression of several stromal- and mesenchymal-associated genes in the combination group, including FAP, PRELP, SCARF2, FHL1 and TSPAN7, as well as the monocyte/macrophage-recruiting chemokine CCL2 (Figure 5D, Supplementary Figure 4). Pathway analysis confirmed the negative enrichment of EMT, G2M checkpoint, E2F targets, MYC targets, angiogenesis and TGFβ signalling (Figure 5E, Supplementary Figure 5C). Consistently, Gene Ontology analysis showed reduced enrichment of collagen-containing extracellular matrix, extracellular structure organisation and cell division-related processes in the combination group (Figure 5F, Supplementary Figure 5D). These findings indicate that adding pembrolizumab to trastuzumab attenuates pathways linked to mesenchymal transition, stromal remodelling and proliferation. Notably, both direct comparisons, combination versus pembrolizumab and combination versus trastuzumab, showed shared strong negative enrichment of G2M checkpoint, E2F targets and MYC targets V1, confirming that the combination consistently suppresses proliferative transcriptional programmes beyond either monotherapy alone. This was further supported by positive enrichment of KRAS signalling down in both comparisons.

These findings suggest that the two agents contribute complementary pathway-level effects: trastuzumab enhances interferon responses, T cell activation and adaptive immune signalling when added to pembrolizumab, whereas pembrolizumab attenuates trastuzumab-associated EMT, TGFβ and ECM-remodelling programmes.

### Combination-associated transcriptional divergence identifies suppression of ECM, EMT and IL6-JAK-STAT3-associated programmes

To identify transcriptional changes preferentially associated with combination treatment, we first compared the log₂ fold-change of each gene in the combination group against the corresponding monotherapy response (Figure 6A, B). Genes above or below the concordance line were interpreted as being more strongly induced or suppressed by the combination than by the respective monotherapy. This analysis showed moderate concordance between the combination and pembrolizumab responses (r = 0.592), and between the combination and trastuzumab responses (r = 0.557), indicating that the combination shared transcriptional features with both monotherapies while also producing distinct gene-level changes.

**Figure 6.**
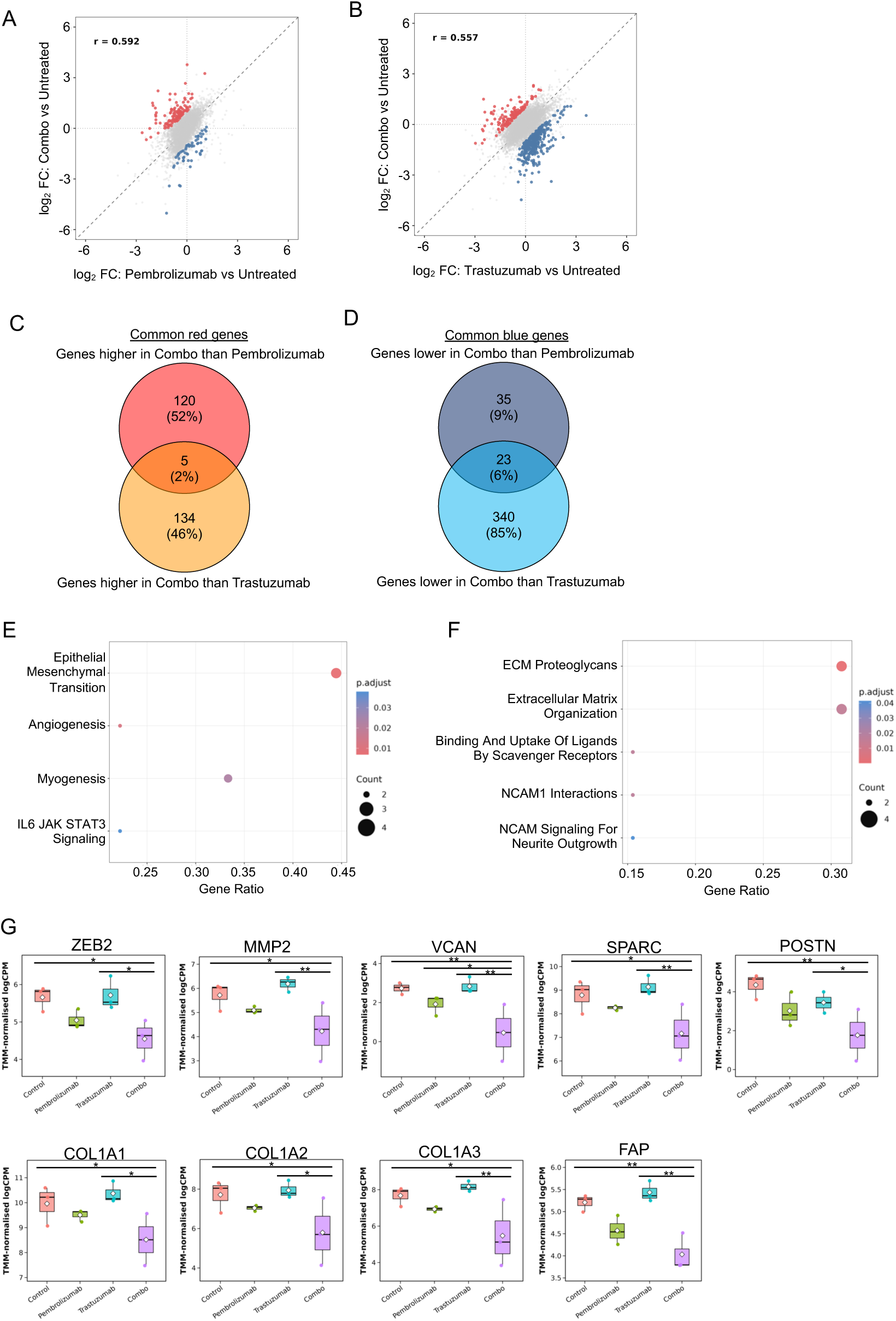
Combination therapy suppresses epithelial-mesenchymal transition and extracellular matrix remodelling relative to monotherapy. Transcriptional effects of combination therapy were compared with those induced by pembrolizumab or trastuzumab monotherapy. (A, B) Concordance plots comparing log2 fold-change values from combination-versus-control with pembrolizumab-versus-control (A) or trastuzumab-versus-control (B), respectively. Genes expressed at higher levels in the combination group than the corresponding monotherapy are shown in red, whereas genes expressed at lower levels are shown in blue. (C, D) Venn diagrams showing genes commonly increased (C) or decreased (D) by combination therapy relative to both monotherapies. (E, F) Hallmark (E) and Reactome (F) enrichment analyses of combination-suppressed gene sets. (G) Expression of indicated mesenchymal, matrix-remodelling and extracellular matrix genes across treatment groups.

We next focused on genes that were consistently altered by the combination relative to both monotherapies. In the upregulated set, 125 genes were higher in the combination group than in pembrolizumab, and 139 genes were higher than in trastuzumab, with only five genes shared between both comparisons: AK8, CYP1B1, OAZ3, TGIF2-RAB5IF and TRAV19 (Figure 6C). In contrast, the downregulated set was larger and more strongly driven by the trastuzumab comparison: 58 genes were lower in the combination group than in pembrolizumab, and 363 genes were lower than in trastuzumab, with 23 genes shared between both comparisons: ADCYAP1R1, ASDURF, CCN2, COL3A1, CXCL14, CXCL3, FILIP1L, HMGN2P5, HOXD8, IL1R2, KCNQ3, LETM2, LINC02009, NCAM1, NR2F1, ROR1, SPARC, TSPAN7, VCAN, VDAC2P5, WBP1P1, WT1 and ZNF423 (Figure 6D).

To define the biological meaning of these combination-suppressed genes, we performed over-representation analysis (ORA) across Hallmark and Reactome gene sets. After filtering for significant pathways and excluding pathways supported by only a single gene, the enriched pathways included EMT, angiogenesis, myogenesis, IL6-JAK-STAT3 signalling, ECM proteoglycans and extracellular matrix organisation, suggesting that combination treatment preferentially attenuated a mesenchymal and matrix-remodelling programme relative to both monotherapies (Figure 6E, F).

These pathway-level findings were further supported by targeted gene-level validation. Combination-treated tumours showed reduced expression of key EMT-associated genes, including ZEB2 and MMP2, as well as extracellular matrix and stromal-remodelling genes such as VCAN, SPARC and POSTN (Figure 6G, H). Structural collagen genes, including COL1A1, COL1A2 and COL1A3, and the activated CAF marker FAP, were also reduced in the combination group. Notably, in most cases, neither pembrolizumab nor trastuzumab monotherapy substantially reduced the expression of these genes.

### Transcriptional programmes preferentially detected in the combination-versus-control group

We next compared the overlap of genes across the three treatment-versus-control groups. Genes in the combination-versus-control group but not in either monotherapy-versus-control comparison were classified as combination-unique genes. This identified 91 upregulated and 308 downregulated genes (Figure 7B).

**Figure 7.**
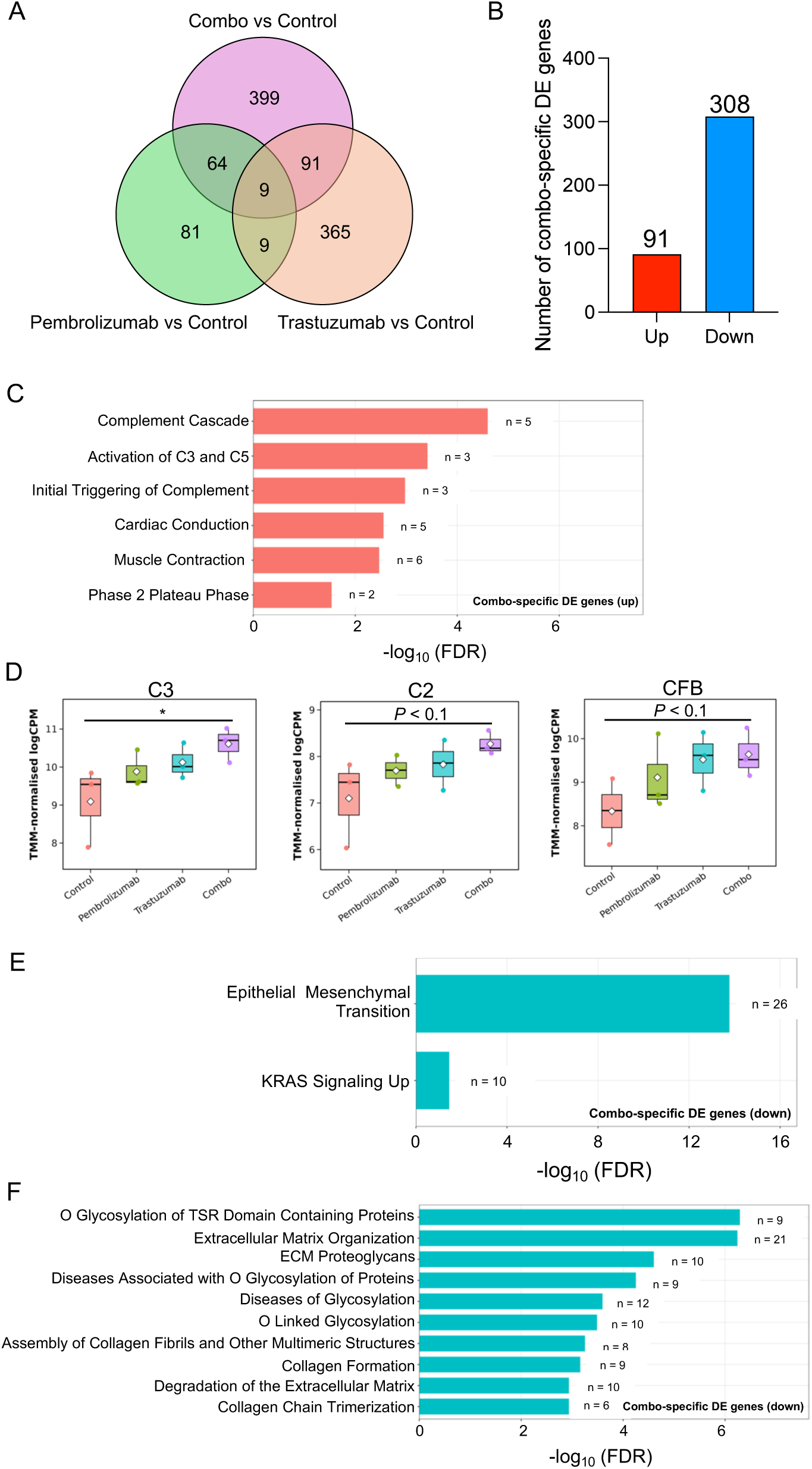
Treatment-associated transcriptional changes preferentially detected in the combination-versus-control comparison. Genes meeting the differential expression threshold in the combination-versus-control comparison were compared with those meeting the same threshold in the pembrolizumab-versus-control and trastuzumab-versus-control comparisons. (A) Venn diagram showing overlap between differentially expressed genes across the three treatment-versus-control comparisons. (B) Number of upregulated and downregulated genes meeting the differential expression threshold only in the combination-versus-control comparison. (C) Reactome enrichment analysis of upregulated genes meeting the differential expression threshold only in the combination-versus-control comparison.. (D) Expression of selected complement-associated genes across treatment groups. (E, F) Hallmark (E) and Reactome (F) enrichment analyses of downregulated genes meeting the differential expression threshold only in the combination-versus-control comparison..

Over-representation analysis of the combination-unique upregulated genes revealed enrichment of complement-related Reactome pathways, including complement cascade and initial triggering of complement (Figure 7C). To explore this further, we examined complement-related genes and identified upregulation of C2, C3 and CFB in the combination group (Figure 7D). Other upstream complement components, including C1R, C1S, C4A and C4B, showed a similar upward trend in the combination group, whereas downstream complement-associated gene C5 were not coordinately increased (Supplementary Figure 6). These findings indicate increased expression of several upstream complement-associated transcripts in the combination group, although functional complement activation remains to be validated.

In the combination-unique downregulated gene set, Hallmark enrichment again identified EMT as the dominant suppressed programme, together with KRAS signalling (Figure 7E). Reactome enrichment further highlighted extracellular matrix organisation, ECM proteoglycans, collagen formation and extracellular matrix degradation.

Overall, these findings suggest that the enhanced anti-tumour activity of pembrolizumab plus trastuzumab is associated with suppression of tumour cell proliferation, EMT and ECM remodelling, rather than a simple addition of the two monotherapy responses. In parallel, the emergence of complement-associated pathways among combination-unique upregulated genes suggests that combination treatment may induce a broader immune-activating complement response than trastuzumab monotherapy alone.

## Discussion

Combination of pembrolizumab, trastuzumab and chemotherapy has improved clinical outcomes in HER2-positive advanced gastric and gastro-oesophageal junction adenocarcinoma^12^. However, the specific contribution of pembrolizumab-trastuzumab crosstalk remains difficult to define clinically, as chemotherapy can remodel the tumour immune microenvironment. Our chemotherapy-free model therefore provides a focused experimental setting in which the biological interaction between PD-1 blockade and HER2-targeted therapy can be examined without the confounding effects of cytotoxic chemotherapy.

We demonstrated that pembrolizumab and trastuzumab monotherapy, respectively, led to distinct transcriptional states. Pembrolizumab alone induced a relatively limited transcriptional response and did not generate broad immune-inflammatory remodelling. In contrast, trastuzumab induced broader transcriptional changes, including enrichment of inflammatory signalling, T cell activation and adaptive immune response pathways. These findings are consistent with the known ability of trastuzumab to engage Fc receptor-bearing immune effector cells and promote immune-mediated tumour cell killing in addition to HER2 receptor blockade^17–19^.

However, trastuzumab monotherapy also induced transcriptional programmes that may limit the durability of its anti-tumour effect. Trastuzumab-treated tumours showed increased EMT, TGFβ signalling, angiogenesis, CAF-associated signatures, ECM programmes, and mixed myeloid and regulatory T cell-associated signatures. This suggests that trastuzumab can initiate anti-tumour immune activation, but that this response may be accompanied by compensatory stromal, mesenchymal and immune-regulatory remodelling, which are closely linked to tumour plasticity, immune exclusion, invasion and therapeutic resistance^20,21^. This mixed immune-activating and tissue-remodelling state may help explain why trastuzumab monotherapy reduced tumour growth but did not achieve the more pronounced tumour control observed with the combination treatment.

One of the most prominent features of the combination treatment was suppression of EMT and ECM remodelling. This is particularly relevant in gastric cancer, where mesenchymal and stromal-rich tumours are strongly associated with aggressive disease biology^22–24^. In the Asian Cancer Research Group molecular classification, the mesenchymal-like MSS/EMT subtype was reported to have the worst prognosis among the four gastric cancer subtypes, the highest recurrence frequency of 63%, and a strong tendency towards peritoneal seeding^25^. This clinical association supports the biological importance of EMT-related programmes in gastric cancer progression and dissemination. Consistent with this, EMT-based transcriptional signatures have been shown to predict prognosis and therapeutic response in gastric cancer^26–29^.

ECM remodelling is also closely connected to this mesenchymal tumour state^30,31^. Gastric cancer studies have shown that ECM-related signatures can stratify patients by prognosis and chemotherapy response, suggesting that the matrix remodelling state is clinically informative rather than a passive stromal feature^32–34^. Proteomic analyses of decellularised human gastric cancer tissues have further demonstrated marked ECM heterogeneity across histological subtypes, with tumour-enriched matrisome-associated proteins and subtype-specific ECM composition^35^. These findings indicate that the ECM is a crucial biological component of gastric cancer progression. Mechanistically, collagen deposition, ECM organisation and CAF-associated matrix remodelling can increase tissue stiffness, promote invasion and angiogenesis, and generate a tumour-supportive niche^36–38^. Recent work has also shown that collagen-rich ECM can promote immune evasion in gastric cancer through IL4I1-AHR signalling, linking matrix remodelling directly to impaired anti-tumour immunity^39^.

In addition to these dominant immune and stromal-remodelling effects, our analysis identified enrichment of complement-related pathways in combination-treated tumours. Increased expression of several upstream complement-associated genes, including C2, C3 and CFB, suggests that complement-related transcriptional activity may represent an additional feature associated with combination treatment. Trastuzumab is known to engage Fc-dependent immune mechanisms, providing a potential biological context for these observations. However, complement protein activation, Fc-dependent effector activity and the contribution of specific immune cell populations were not directly assessed in the present study. These findings should therefore be regarded as hypothesis-generating rather than evidence of functional complement activation or a defined Fc-mediated mechanism.

This study has several limitations. The PBMC-engrafted humanised mouse model allows assessment of human immune-tumour interactions *in vivo*, but it does not fully reproduce the complexity of the human gastric cancer immune microenvironment. This model also carries a potential risk of graft-versus-host disease (GvHD). Therefore, all experiments were completed within four weeks since PBMC engraftment, when GvHD-related effects are expected to be minimal. In addition, although the absence of chemotherapy was intentional and enabled the pembrolizumab-trastuzumab interaction to be elucidated, it also means that the findings do not directly model the approved first-line clinical regimen. In addition, bulk RNA-seq captures integrated the transcriptional states of tumour cells and the overall microenvironment, and cannot assign pathway changes to specific cell populations. Collectively, the mechanistic insights inferred from the transcriptomic findings require validation at the protein level and in additional models.

In summary, our findings provide transcriptional insight into how pembrolizumab and trastuzumab remodel the tumour microenvironment of gastric cancer. Combination treatment was associated with enhanced cytotoxic immune signatures and reduced proliferative, EMT- and ECM-remodelling transcriptional programmes..

## Materials and Methods

### Animals and housing conditions

Six- to eight-week-old female NOG mice were obtained from Beijing Vital River Laboratory Animal Technology Co., Ltd. Mice were maintained under specific pathogen-free conditions in a barrier facility at Shanghai Qishang Biomedical Technology Co., Ltd. Animals were housed under controlled environmental conditions, with a temperature of 20-26°C, relative humidity of 40-70%, and a 12 h light/12 h dark cycle.

### Animal model

The human gastric cancer cell line NCI-N87 was obtained from ATCC and cultured in complete RPMI-1640 medium. To establish PBMC-humanised tumour-bearing mice, NOG mice received 0.2 mL of PBMC suspension, corresponding to 3 x 10⁶ PBMCs per mouse, by tail-vein injection on day 0. Three days after PBMC transfer, mice were implanted subcutaneously with NCI-N87 cells into the right anterior dorsal flank. Each mouse received 0.1 mL of cell suspension containing 10 x 10⁶ NCI-N87 cells in 30% Matrigel basement membrane matrix. Two days after tumour implantation, mice were randomised into four groups, with six mice per group: vehicle control, pembrolizumab, trastuzumab, and pembrolizumab plus trastuzumab. Mice received PBS as vehicle control, pembrolizumab at 10 mg/kg by tail-vein injection, trastuzumab at 4 mg/kg by subcutaneous injection, or the corresponding combination at the same doses and routes. Treatments were administered twice weekly for three consecutive weeks.

Tumour volume were recorded twice weekly throughout the treatment period. Tumour length and width were measured using digital callipers, and tumour volume was calculated as: volume = (length x width²)/2. At the experimental endpoint, mice were euthanised by intraperitoneal injection of an overdose of sodium pentobarbital (150 mg/kg, prepared in sterile saline). Following complete loss of the corneal reflex and spontaneous respiration, cervical dislocation was performed as a secondary method to confirm death. Tumours were then excised, measured, weighed, and collected for downstream transcriptomic analysis.

### RNA extraction and bulk RNA sequencing

Total RNA was extracted from tumour tissues using TRIzol Reagent (Invitrogen, USA) according to the manufacturer’s protocol. RNA concentration and purity were assessed using an ND-2000 spectrophotometer (NanoDrop Technologies, USA), and RNA integrity was evaluated using a 2100 Bioanalyser (Agilent, USA). Samples meeting the predefined quality-control criteria (OD260/280 = 1.8-2.2, OD260/230 ≥ 2.0, RIN ≥ 6.5, 28S:18S ≥ 1.0, and total RNA amount >2 μg) were used for library preparation. Stranded mRNA libraries were generated using the Illumina Stranded mRNA Prep, Ligation kit (Illumina, USA) and sequenced on a Novaseq 6000 sequencer (Illumina, USA).

### Bulk RNA-sequencing data processing

Bulk RNA-sequencing was performed on tumour tissues from 12 samples, comprising vehicle control, pembrolizumab, trastuzumab, and pembrolizumab plus trastuzumab groups, with three biological replicates per group. Sequencing and primary processing were performed by Majorbio. Raw sequencing reads were quality-filtered to generate clean reads and aligned to the human GRCh38 reference genome, followed by gene-level read quantification. The resulting annotated gene-level count matrix, containing Ensembl gene identifiers, gene symbols, gene length information and sample-level read counts, was used as the starting point for all downstream analyses.

### Normalisation and differential gene-expression analysis

Downstream analysis was performed in R using the annotated gene-level count matrix. Prior to normalisation and differential expression testing, lowly expressed genes were removed using “filterByExpr” function from ‘edgeR’. Library-size normalisation was then performed using the trimmed mean of M-values method, and normalised log2 counts per million values were generated for exploratory analysis and visualisation.

Differential gene-expression analysis was performed using the limma-voom framework from ‘limma’ package. The comparisons were pembrolizumab versus control, trastuzumab versus control, combination treatment versus control, combination treatment versus pembrolizumab, and combination treatment versus trastuzumab. Volcano plots were generated using log2 fold-change and -log10 P value. Heatmaps were generated from TMM-normalised logCPM values.

### Gene set enrichment analysis

Gene set enrichment analysis was performed using the ‘fgsea’ package. Genes were ranked using the limma moderated t-statistic. Gene sets were obtained from the Molecular Signatures Database using ‘msigdbr’, including Hallmark and C1-C8 collections. Gene sets containing fewer than 10 or more than 500 detected genes were excluded.

### Single-sample gene set enrichment analysis

Single-sample gene set enrichment analysis was performed using “GSVA” on TMM-normalised logCPM values. Hallmark gene sets were combined with curated immune-, tumour-, and stromal-related signatures. Signature scores were calculated for each sample and z-scored across samples for heatmap visualisation. Group-level signature activity was summarised by averaging individual-sample ssGSEA scores within each treatment group.

### xCell immune deconvolution

Immune cell-state enrichment scores were estimated using the xCell method implemented in the ‘immunedeconv’ R package. Raw counts from the annotated count matrix were normalised by gene length and library size to generate a TPM-normalised gene-symbol expression matrix. xCell scores were visualised after row-wise z-score transformation to compare inferred immune cell-state signatures across treatment groups.

### Ethics statement

All animal procedures were performed in accordance with institutional animal welfare requirements and were approved by the Animal Welfare and Ethics Committee of Shanghai Qishang Biomedical Technology Co., Ltd. (approval number: IACUC-RL-01).

### Statistical analysis

Longitudinal tumour-growth curves were analysed using two-way ANOVA followed by Tukey’s post hoc multiple comparisons test. Area under the tumour-growth curve was calculated from mean tumour-volume values over time and used to assess overall treatment effect. Statistical significance was denoted in figures as follows: \**P* < 0.05; \*\**P* < 0.01; \*\*\**P* < 0.001; \*\*\*\**P* < 0.0001; NS: not significant.

## Availability of data and materials

The RNA-seq data generated from this study can be obtained from the NCBI GEO database with accession number GSE335301. All data needed to evaluate the conclusions in this paper are present in the paper and the supplementary materials.

## Competing interests

The authors declare no competing interests.

## Fundings

2023 Chongqing Municipal Education Commission Science and Technology Research Program Project (#KJZD-M202300202)

2024 Horizontal Research Project (#F2024126)

## Authors’ contributions

Conceptualisation: PKTW, JW;

Methodology: PKTW, JW;

Data acquisition: SP, XZ;

Data analysis: PKTW, JW;

Manuscript writing: PKTW.

## Acknowledgments

Not applicable

## Supplementary Figures

**Supplementary Figure 1.**
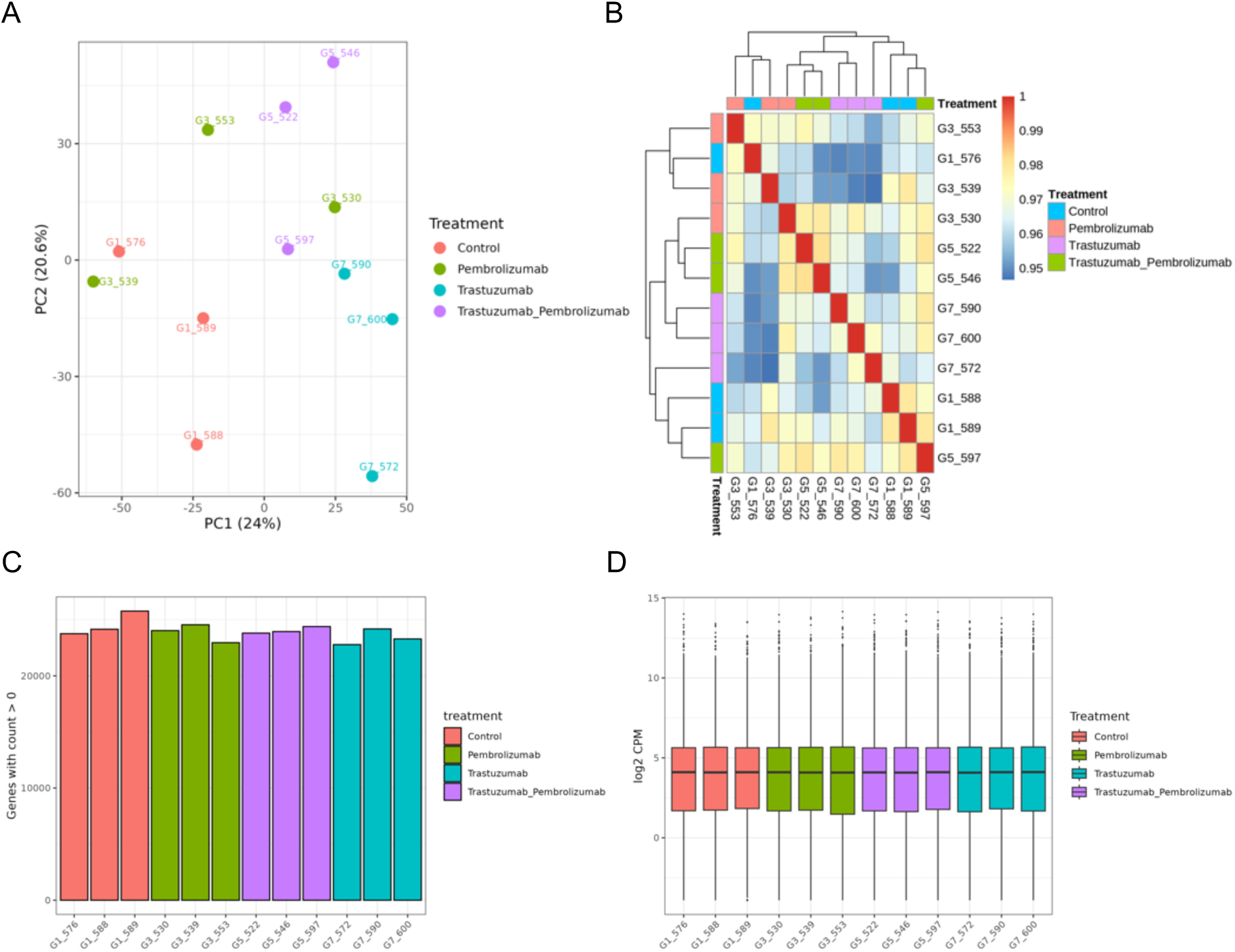
Quality control analysis of bulk RNA-seq samples. Principal component analysis (A), sample correlation (B), detected-gene counts (C) and logCPM distribution (D) plots were generated from the filtered, TMM-normalised expression matrix. Samples are coloured by treatment group.

**Supplementary Figure 2.**
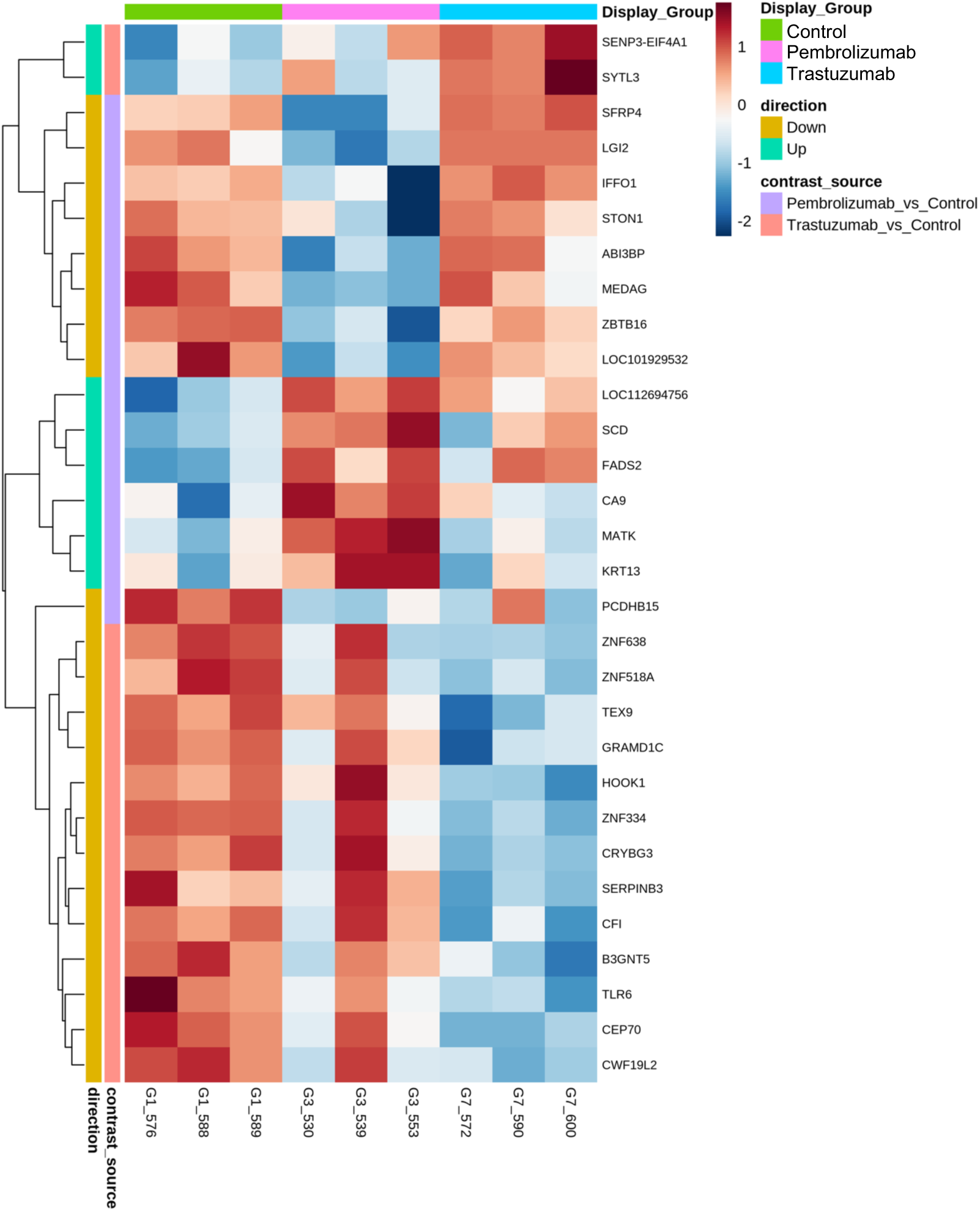
Top differentially expressed genes following monotherapy treatment. Heatmap showing top differentially expressed genes from the pembrolizumab-versus-control and trastuzumab-versus-control comparisons. Columns represent individual tumour samples and are annotated by treatment group.

**Supplementary Figure 3.**
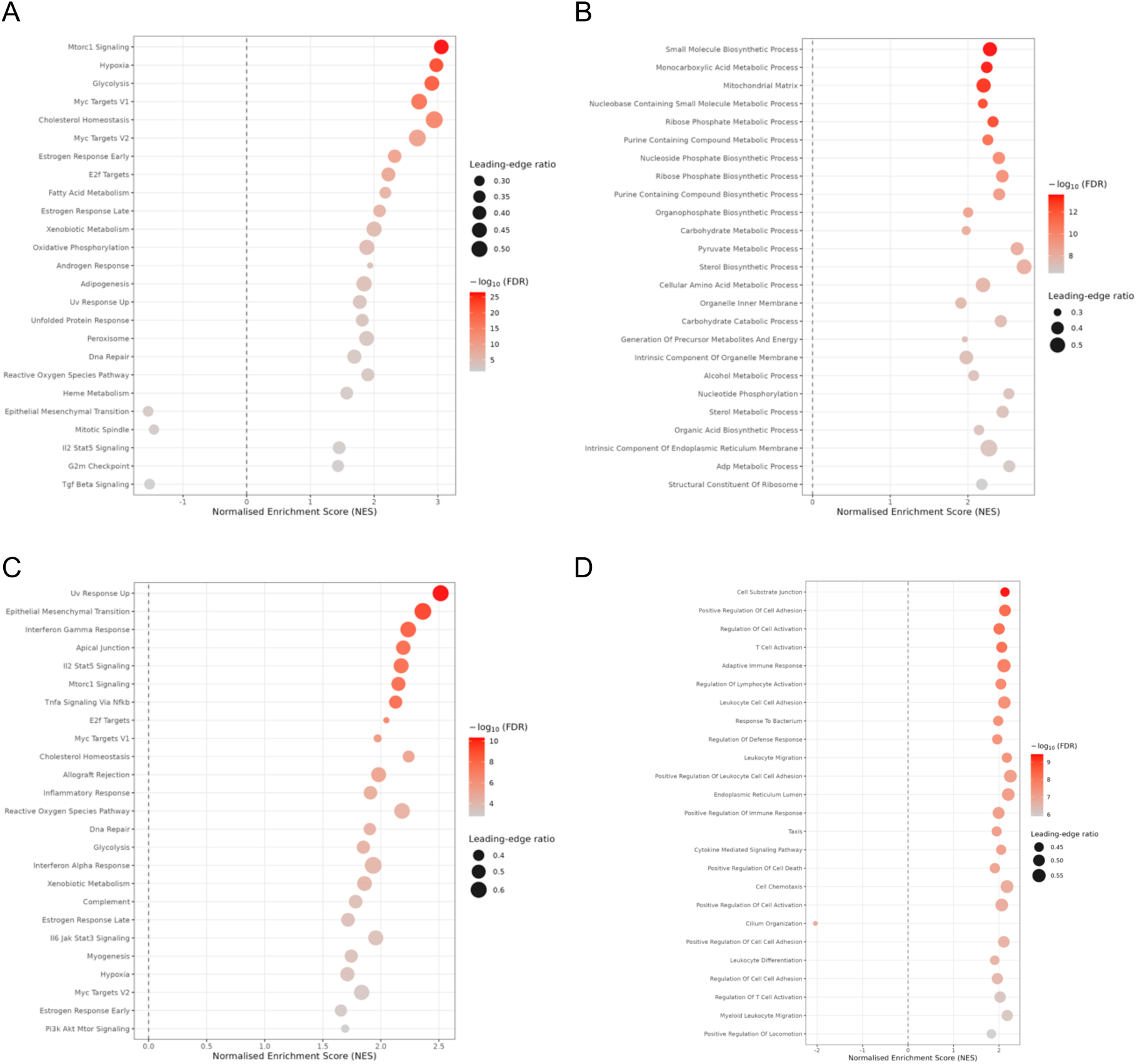
GSEA of monotherapy versus control comparisons. Dot plots showing the top enriched pathways identified by Hallmark and C5 Gene Ontology GSEA. Hallmark pathways (A) and C5 Gene Ontology pathways (B) are shown for pembrolizumab versus control, while Hallmark pathways (C) and C5 Gene Ontology pathways (D) are shown for trastuzumab versus control. Dot colour represents -log10(FDR), and dot size represents the leading-edge ratio. Positive NES indicates enrichment in the corresponding monotherapy group, whereas negative NES indicates enrichment in the control group.

**Supplementary Figure 4.**
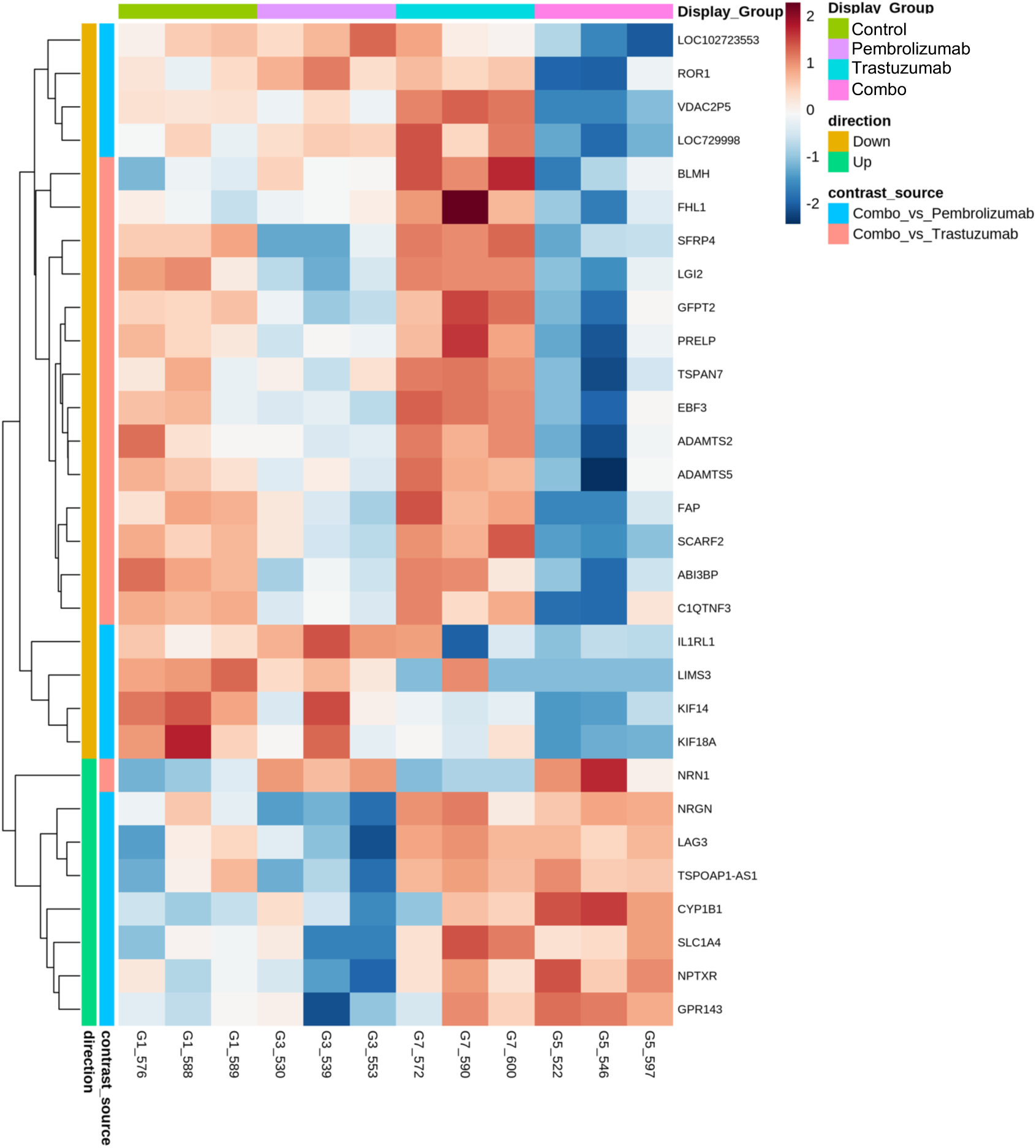
Differentially expressed genes in combination versus monotherapy comparisons. Heatmap showing top differentially expressed genes from the combination-versus-pembrolizumab and combination-versus-trastuzumab comparisons. Columns represent individual tumour samples and are annotated by treatment group.

**Supplementary Figure 5.**
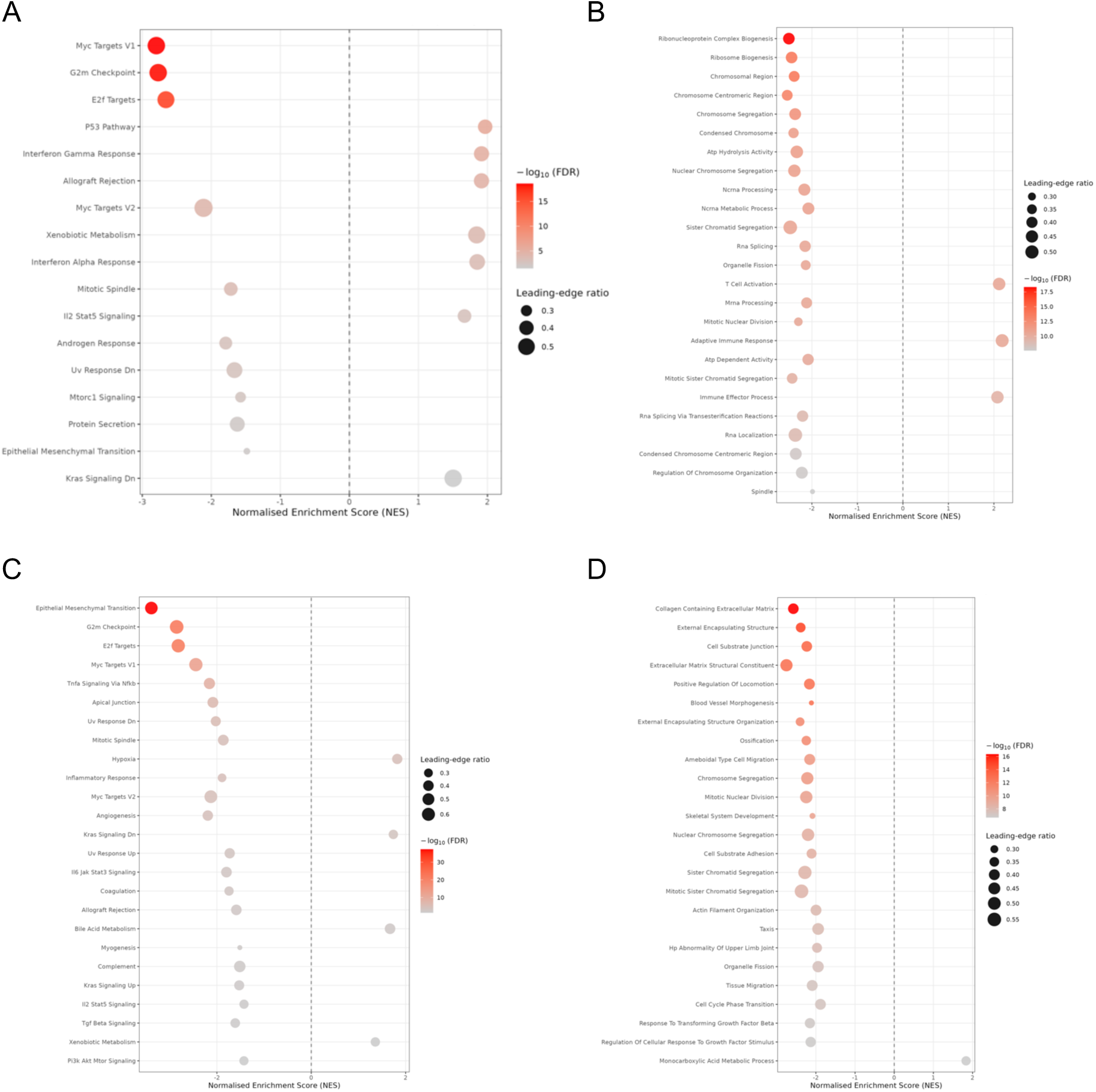
GSEA of combination versus monotherapy comparisons. Dot plots showing the top enriched pathways from Hallmark and C5 Gene Ontology GSEA. Hallmark pathways (A) and C5 Gene Ontology pathways (B) are shown for combination versus pembrolizumab. Hallmark pathways (C) and C5 Gene Ontology pathways (D) are shown for combination versus trastuzumab. Dot colour represents -log10(FDR), and dot size represents the leading-edge ratio. Positive NES indicates enrichment in the combination group, whereas negative NES indicates enrichment in the corresponding monotherapy group.

**Supplementary Figure 6.**
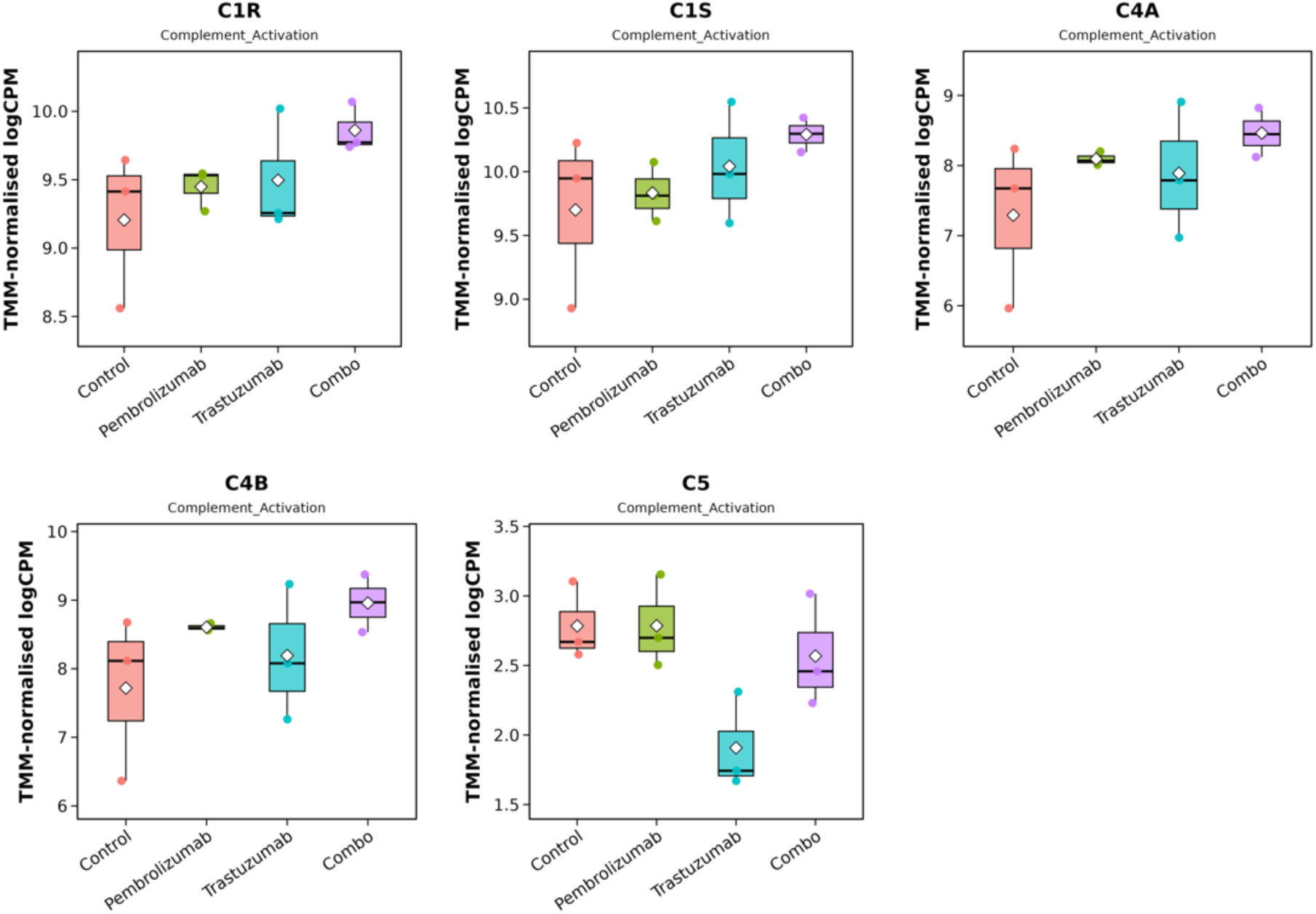
Expression of complement activation genes across treatment groups. Boxplots show TMM-normalised logCPM expression of C1R, C1S, C4A, C4B and C5 across control, pembrolizumab, trastuzumab and combination treatment groups. Each point represents an individual sample, with diamonds indicating group means.

## Notes

### Competing Interest Statement

The authors have declared no competing interest.

